# Comparative Modelling of Actin–Tropomyosin Interfaces

**DOI:** 10.64898/2026.07.06.736648

**Authors:** Revathy Menon, Mohan Balasubramanian, Ramanathan Sowdhamini

## Abstract

Tropomyosins are coiled-coil dimers that polymerize head-to-tail along actin filaments. They stabilize distinct filament populations and regulate the access of myosins and actin-binding proteins in both muscle and non-muscle contexts. Despite their central regulatory role, how filament length and isoform identity of different tropomyosin homologues might modulate actin affinity is not completely understood, especially across species. Here, we present a stepwise computational docking pipeline combining AlphaFold2-Multimer coiled-coil models, experimentally informed residue-level restraints, and pseudo-energy analysis *via* PPCheck to build and evaluate actin-tropomyosin co-polymer models for three isoforms: human TPM1 (hTPM1; 284 residues), human TPM4 (hTPM4; 248 residues), and *Schizosaccharomyces pombe* Cdc8 (SpCdc8; 161 residues). Interface energetics reveal a consistent hierarchy in which the shortest filament, SpCdc8, achieves the most stabilizing and residue-rich actin contacts, consistent with reduced cumulative geometric penalty along the actin helix. Among human isoforms, hTPM1 forms stronger interfaces with actin than hTPM4. The hTPM1-actin model also exhibits higher contact density and additional energetic hotspots, in agreement with the experimentally established slower exchange kinetics of TPM1 isoforms on actin filaments relative to TPM4. Hotspot mapping identifies conserved acidic residues at equivalent positions across all three isoforms, emphasizing the importance of electrostatic anchor points in maintaining interface integrity across diverse evolutionary contexts. Modeling of four temperature-sensitive SpCdc8 mutations (A18T, R21H, E31K and E129K) reveals that these substitutions substantially destabilize the coiled-coil dimer without significantly affecting actin interactions, suggesting that subtle regulatory failure arises from compromised longitudinal cable continuity rather than from direct loss of actin affinity. Taken together, our results support a hierarchical model of tropomyosin dimer stability, actin–tropomyosin recognition in which filament length imposes a geometric baseline on interface stability, onto which isoform-specific sequence evolution superimposes functional tuning. The tropomyosin homologues we studied appear to retain conserved electrostatic hotspots thereby providing a common structural scaffold across tissues and organisms.

## Introduction

The actin cytoskeleton underpins a wide range of cellular functions, from sarcomeric contraction in muscle to cell shape regulation, division, migration and intracellular trafficking in non-muscle cells. Its versatility arises from the ability of actin filaments to form biochemically distinct populations that are selectively decorated by different actin-binding proteins (ABPs). Tropomyosins are central to this specialization: they are α-helical coiled-coil dimers that polymerize head-to-tail along the major groove of F-actin, stabilizing the filament and regulating the access of myosins and numerous ABPs to specific subsets of filaments in both muscle and non-muscle contexts (Gunning et al., 2008; Jansen and Goode, 2019; Vindin and Gunning, 2013). In striated muscle, tropomyosin works together with the troponin complex to confer Ca²⁺-dependent switching between blocked, closed and open states of the thin filament, whereas in the cytoskeleton, distinct tropomyosin isoforms partition to different actin networks and tune their mechanical and signaling properties in a largely Ca²⁺-independent manner (Gunning et al., 2015; Narita et al., 2001; Rynkiewicz et al., 2022).

Despite this central regulatory role and the extreme sequence conservation of the actin–tropomyosin binding interface across eukaryotes, there remain important gaps in our structural understanding of the acto–tropomyosin copolymer. From an evolutionary perspective, this is particularly striking given that tropomyosin itself varies substantially in length and sequence, ranging from long, muscle-associated isoforms in vertebrates to shorter cytoskeletal and fungal isoforms. Early three-dimensional reconstructions of thin filaments relied on relatively low-resolution cryo-EM and often used truncated or segmented constructs, which limited insight into how a continuous tropomyosin cable tracks the helical actin lattice (Behrmann et al., 2012; Sousa et al., 2013). Subsequent higher-resolution structures of F-actin–tropomyosin complexes and rigor actin–tropomyosin–myosin assemblies have clarified average tropomyosin positions and regulatory states (von der Ecken et al., 2016), but they primarily focus on muscle isoforms and frequently employ constructs that sample only part of the full regulatory repeat (Doran et al., 2023; von der Ecken et al., 2015). Recent single-molecule imaging (Cagigas et al., 2025) and modeling studies reinforce the view that tropomyosin forms a continuous, head-to-tail polymer along actin and that isoforms can follow distinct paths and impose isoform-specific regulation (Selvaraj et al., 2023; Siatkowska et al., 2025).

Advances in computational structural biology now offer powerful routes to bridge this structural gap. AlphaFold2 (Jumper et al., 2021) has demonstrated remarkable accuracy in modeling coiled-coil domains, capturing both local geometry and global topology across dimeric and higher-order coiled-coil assemblies (Madaj et al., 2025). More recently, AlphaFold3 (Abramson et al., 2024) extended this paradigm to directly predict joint structures of protein complexes and other biomolecular assemblies using a diffusion-based architecture, substantially improving the accuracy of complex modeling relative to earlier methods (Krokidis et al., 2025). In parallel, integrative modeling approaches that combine structural prediction, cryo-EM density and biophysical restraints have been used to build thin-filament models and to map energetic hotspots on tropomyosin, highlighting the importance of specific quasi-repeats (notably Period 2 and related segments) as key contributors to actin affinity and cooperative activation (Margaret Sunitha et al., 2012; Pavadai et al., 2020). However, much of this structural knowledge is derived from human or vertebrate systems, and comparatively little is known about how differences in tropomyosin length and sequence across organisms influence actin binding and filament mechanics. These developments provide a framework in which high-confidence coiled-coil models can be docked onto F-actin using experimental restraints, yielding testable hypotheses about isoform-specific interfaces and regulatory mechanisms.

Fission yeast (*Schizosaccharomyces pombe)* tropomyosin Cdc8 provides a complementary system for dissecting thin-filament mechanics. Cdc8 is essential for contractile ring assembly and actin cable organization, and temperature-sensitive (ts) cdc8 alleles have long been used to probe how single-residue substitutions perturb filament stability and function (Balasubramanian et al., 1992; Johnson et al., 2018; Rajagopalan et al., 2004). Recent biochemical and structural work has begun to clarify how N-terminal acetylation and specific mutations alter Cdc8 cable formation, actin binding and thermal stability, but a unified, atomistic explanation of how these changes impact the actin–tropomyosin copolymer is still emerging (Reinke et al., 2024; Tang et al., 2023). Notably, both yeast Cdc8 and human TPM4 are shorter than canonical muscle tropomyosins, raising the question of how filament length itself might modulate actin affinity and stability.

Here, we have used a state-of-the-art and stepwise computational modeling pipeline to address these outstanding questions. We first generate full-length coiled-coil models of human TPM1, human TPM4 and *S. pombe* Cdc8 (referred to as hTPM1, hTPM4, and SpCdc8 respectively throughout the manuscript for brevity). We dock G-actin consecutively onto them, using experimentally informed restraints, to generate one actin proto-filament. We then quantify how tropomyosin filament length and isoform identity modulate the energetics and residue-level architecture of the actin interface, using pseudo-energies. Finally, we model a set of SpCdc8 temperature-sensitivity-inducing mutations and evaluate their impact, providing a structural framework for understanding how subtle sequence changes can destabilize the actin–tropomyosin copolymer and how mutations in tropomyosin compromise function.

## Materials and Methods

### Generation and assessment of AlphaFold models for human tropomyosins, yeast CDC8, and yeast actin proteins

The sequences of the human TPM1 and TPM4 proteins were obtained from the UniProt entries P09493 and P67936, respectively. The human TPM1 protein is 284 amino acids long, while the TPM4 protein is 248 residues long. The sequence of the CDC8 protein from *Schizosaccharomyces pombe* was obtained from the UniProt entry Q02088. The sequence is 161 amino acids long, and the DeepCoil2 and CoCoNat servers were used to validate the coiled-coil structure of the molecule.

As full-length structures of these proteins were not available from the Protein Data Bank, AlphaFold2-Multimer (AF2; via ColabFold) was used to model the tropomyosin and CDC8 filaments. AF2 was chosen here because it is well validated for coiled-coil systems and reliably captures parallel dimeric architecture and heptad register across extended α-helical assemblies (Madaj et al., 2025). These models were relaxed, and all other parameters were set to default. Each AlphaFold2 run yielded top five ranked models. While we accounted for the rank assigned by AlphaFold2, we also utilised PPCheck - an in-house in-house web server capable of identifying interactions between two interacting proteins, calculating pseudoenergies for pairs of interacting interfaces, identifying hotspots at the interface, and predicting near-native poses from a set of docked decoys. We compared the PPCheck energies of these models, and selected the model with the most stabilizing PPCheck energies for docking.

Because the structure of actin from *S. pombe* is also not available in the Protein Data Bank, AlphaFold3 (AF3) was used to model this molecule, with sequence data from the UniProt entry P10989. All modelling parameters were set to default. The sequence is 375 amino acids long. AF3 was selected in this case because it has been shown to perform robustly for globular monomeric proteins and protein–protein contexts beyond coiled-coil systems (Krokidis et al., 2025). As PPCheck quantifies interaction energies at interfaces, we could not use it to identify the top model. All five models were virtually identical, with the same pTM and ranking scores (pTM = 0.92; ranking score = 0.92, PAE = 0.76). Given this near-identity, model 0 was selected as a representative structure for S. pombe actin.

We also assessed the feasibility of full-assembly prediction with AF3 (all parameters set to default) to generate models of the entire macromolecular complex, comprising two strands of tropomyosin/CDC8 with the appropriate number of actins (forming a single filament), but these predictions suffered from poor convergence, inconsistent filament registry, and steric clashes between neighboring actin subunits, and therefore did not reach acceptable quality for downstream analysis. The structure for human actin (PDB ID: 8GSU) was obtained from the Protein Data Bank.

### Docking protocol

Actin–tropomyosin docking was performed at stoichiometries defined by tropomyosin length: human TPM1 (hTPM1; 284 residues) with seven actin subunits, human TPM4 (hTPM4; 248 residues) with six actin subunits, and *S. pombe* CDC8 (SpCdc8; 161 residues) with four actin subunits.

To guide sampling, we derived residue-level restraints from published hTPM1 actin-binding regions and observed equivalences in hTPM4 and CDC8 by ClustalW sequence alignment (Madeira et al., 2024). The sequence alignments included known homologues across organisms, as well as human tropomyosin isoforms. Docking was carried out with LightDock (Roel-Touris et al., 2020), which uses global swarm optimization to generate decoys (10 decoys per swarm; 1,000 steps; all other parameters default). In the initial round, a single actin was docked onto each tropomyosin coiled-coil, decoys were clustered using LightDock’s intra-swarm method, and cluster representatives were evaluated with our in-house tool PPCheck (Sukhwal and Sowdhamini, 2015, 2013) to identify stable tropomyosin-actin interfaces. Representative models were selected based on PPCheck energy and satisfaction of the applied restraints. The selected model then served as the receptor for the next actin, and this restrained docking–cluster–evaluate–select cycle was iterated until the target filament stoichiometry was reached.

### Interface analysis

The interaction interface of each decoy in a docking run was analyzed to determine interaction energies and hotspots utilizing PPCheck. The PPCheck algorithm has been trained on mutations from the Alanine Scanning Energetics Database (Thorn and Bogan, 2001) and validated on mutations from the Binding Interface Database (Fischer et al., 2003). The top nine residues exhibiting the highest degree of interactions, and normalized interaction energy < -1, are designated as hotspots.

### Modelling CDC8 temperature-sensitive mutants

Four known temperature-sensitivity-inducing mutations - A18T, R21H, E31K and E129K - were modelled using the final docked SpCdc8-actin model. Each mutation was incorporated symmetrically on both SpCdc8 chains. The mutant structures were energy-minimized using FoldX (Delgado et al., 2025, 2019) and analyzed using PPCheck.

## Results

### AlphaFold model validation

AlphaFold2 runs for hTPM1, hTPM4 and SpCdc8 yielded top five ranked models (shown in Supplementary Figure S1). The selection of AlphaFold models for tropomyosin was based on parameter values obtained from PPCheck and AlphaFold; details of the same are provided in Supplementary Table S1. For hTPM1, the AF2 model with the highest pLDDT score (88.4) was ranked second by the AF2 method; nevertheless, it exhibited the greatest predicted stability based on PPCheck interaction energies for the coiled-coil interface, and was therefore selected as the receptor for consecutive docking. For hTPM4, the top-ranked AF2 model achieved the highest pLDDT score (90.1) and showed the most favourable PPCheck energies; this dimer was likewise chosen for consecutive docking. In the case of *S. pombe* CDC8, the sequence was first validated as a parallel coiled coil with defined heptad registers using CocoNat and DeepCoil2 (Supplementary Figure S2A). Subsequent AF2 modeling yielded five candidates; the top-ranked model displayed the highest pLDDT score (92.4) but maximum stabilizing energies were observed for the model ranked fifth, which was then selected as the receptor for consecutive docking.

### Docking results

Consecutive docking was carried out with LightDock. For hTPM1, residue-level restraints were derived from previously reported tropomyosin–actin contacts (Table 1). ClustalW alignment was used to identify equivalent actin-binding regions for hTPM4 and SpCdc8 (Supplementary Figure S2B). Analysis of the alignments showed that for hTPM4, the first actin-binding region (P1) corresponds to a mid-point of the first and second actin-binding regions (P1 and P2) of hTPM1, while actin-binding regions P2-6 of hTPM4 correspond directly to P3-7 of hTPM1. SpCdc8 P1-3 corresponds directly to P1-3 of hTPM1, and SpCdc8 P4 is equivalent to hTPM1 P7. These equivalences are represented schematically in Figure 1A. This information was then used to derive equivalent restraints applied in the respective docking runs (Figure 1B). The final structures obtained from consecutive docking are shown in Figure 2.

**Figure 1.**
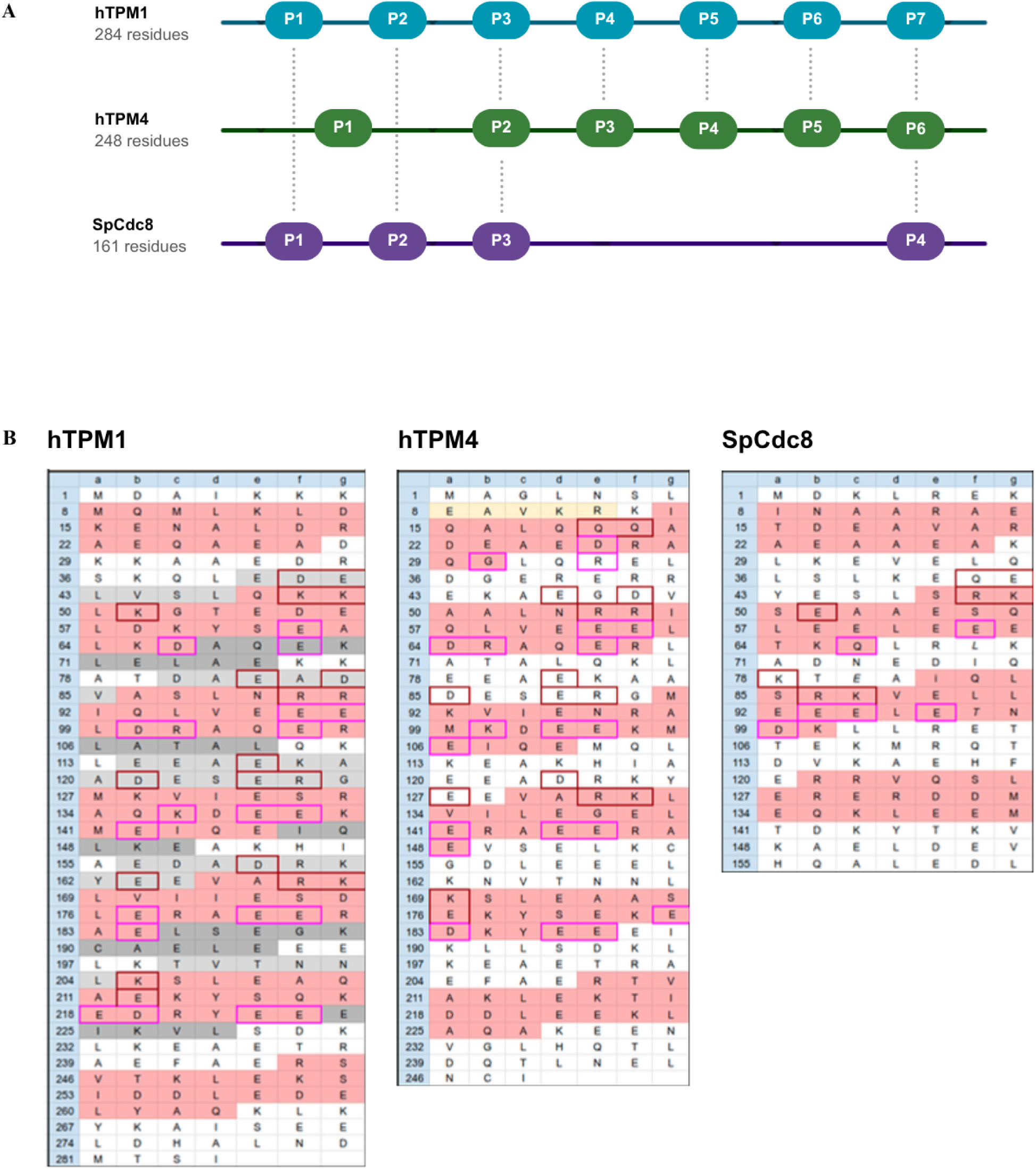
(A) A schematic representing the hTPM1-actin, hTPM4-actin and SpCdc8-actin complexes. The horizontal lines depict the length of the tropomyosin molecules, and the shapes labelled P1-7 represent actin-binding regions on the molecule. The dotted lines are drawn connecting equivalent actin-binding regions. Please note that the diagram is not to scale. (B) The sequences of hTPM1, hTPM4, and pCDC8 split into heptads, with residues labelled a-g per heptad. Pink coloured cells represent actin binding regions adapted from <u>Margaret</u> et al. 2012, and cells with pink and maroon borders highlight key residues. The residue number of each ‘a’ residue is shown on the left side.

**Figure 2.**
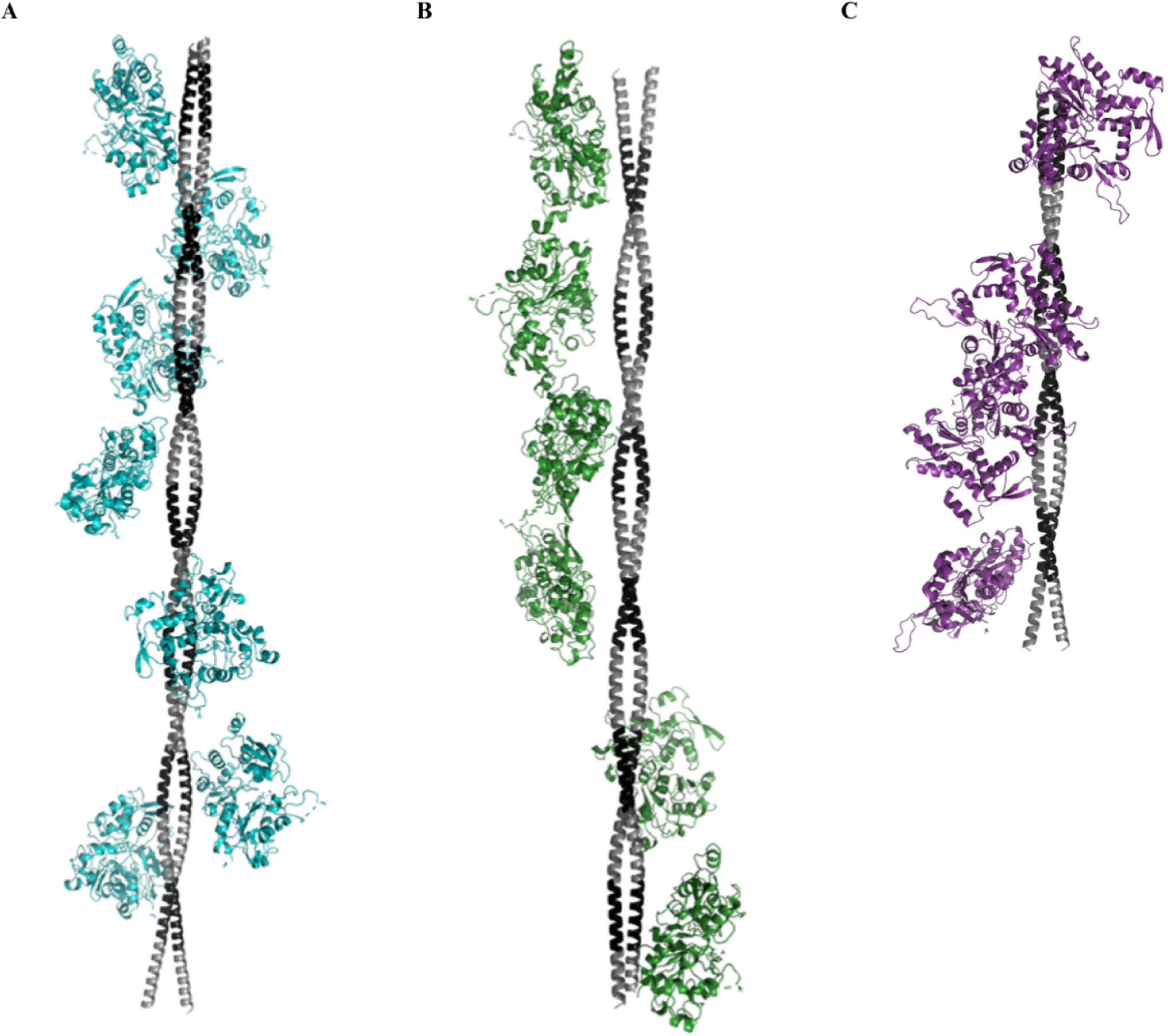
Final structures obtained after consecutive docking for (A) hTPM1-actin, (B) hTPM4-actin and (C) SpCdc8-actin. The coiled-coil filaments are coloured in gray, with the actin-binding regions marked in black.

**Table 1.** Table listing the restraints used for consecutive docking for hTPM1. Data from (Li et al., 2011) and (Sunitha et al., 2012). Equivalent restraints for hTPM4 and SpCdc8 were derived from this table, using the sequence alignment.

| hTPM1 | Actin | Interface |
| --- | --- | --- |
| D258 | R147 | actin subunit 1 - TPM repeat 7 |
| R244 | D25 |  |
| E223 | K326 | actin subunit 2 - TPM repeat 6 |
| D219 | R147 |  |
| E223 | K328 |  |
| D219 | K328 |  |
| E181 | K326 | actin subunit 3 - TPM repeat 5 |
| E181 | R147 |  |
| E181 | K328 |  |
| E184 | K328 |  |
| V170 | P333 |  |
| R167 | D25 |  |
| E142 | K326 | actin subunit 4 - TPM repeat 4 |
| E139 | R147 |  |
| E139 | K328 |  |
| S132 | P333 |  |
| K128 | E334 |  |
| E124 | D25 |  |
| E104 | K326 | actin subunit 5 - TPM repeat 3 |
| E104 | K328 |  |
| D100 | K328 |  |
| E97 | R147 |  |
| R90 | D25 |  |
| E62 | R147 | actin subunit 6 - TPM repeat 2 |
| D58 | R147 |  |
| D58 | K328 |  |
| E62 | K328 |  |
| K48 | D25 |  |
| D41 | R28 |  |
| D20 | R147 | actin subunit 7 - TPM repeat 1 |
| Q9 | D25 |  |

### Energy vs length

Analysis of stabilizing energies (Figure 3; Supplementary Table S2A) revealed a consistent ranking in which CDC8 forms the most stabilizing and residue-rich actin contacts, TPM1 is intermediate and TPM4 is weakest. The most favorable interaction energies correspond to side-on binding of tropomyosin along the major groove of F-actin rather than a superhelical arrangement, indicating that the observed energetic trends reflect physically meaningful filament registry rather than nonspecific surface association. Across all interfaces in the docked complexes, CDC8 displays the most negative normalized stabilizing energies per residue, both within the tropomyosin coiled coil (A–B) and at actin-facing interfaces such as B–E, indicating a generally tighter interface. The shortest filament, CDC8, therefore exhibits the most stabilizing actin–tropomyosin interfaces, consistent with reduced cumulative steric clash along the helical track and a more favorable registry of filament termini within the actin subdomain-1/3 grooves. Within the human isoforms, the A–B interaction remains among the strongest contacts, but TPM1 consistently forms more stabilizing actin-facing interfaces than the slightly shorter TPM4: TPM1 engages more actin-contacting residues per subunit and harbours additional energetic hotspots within canonical actin-binding clusters, whereas several TPM4 interfaces approach neutral energies. The agreement between mean and median per-interface energies (Supplementary Table S2B) indicates that these trends reflect a global shift in interface quality rather than being driven by a few outlier contacts.

**Figure 3.**
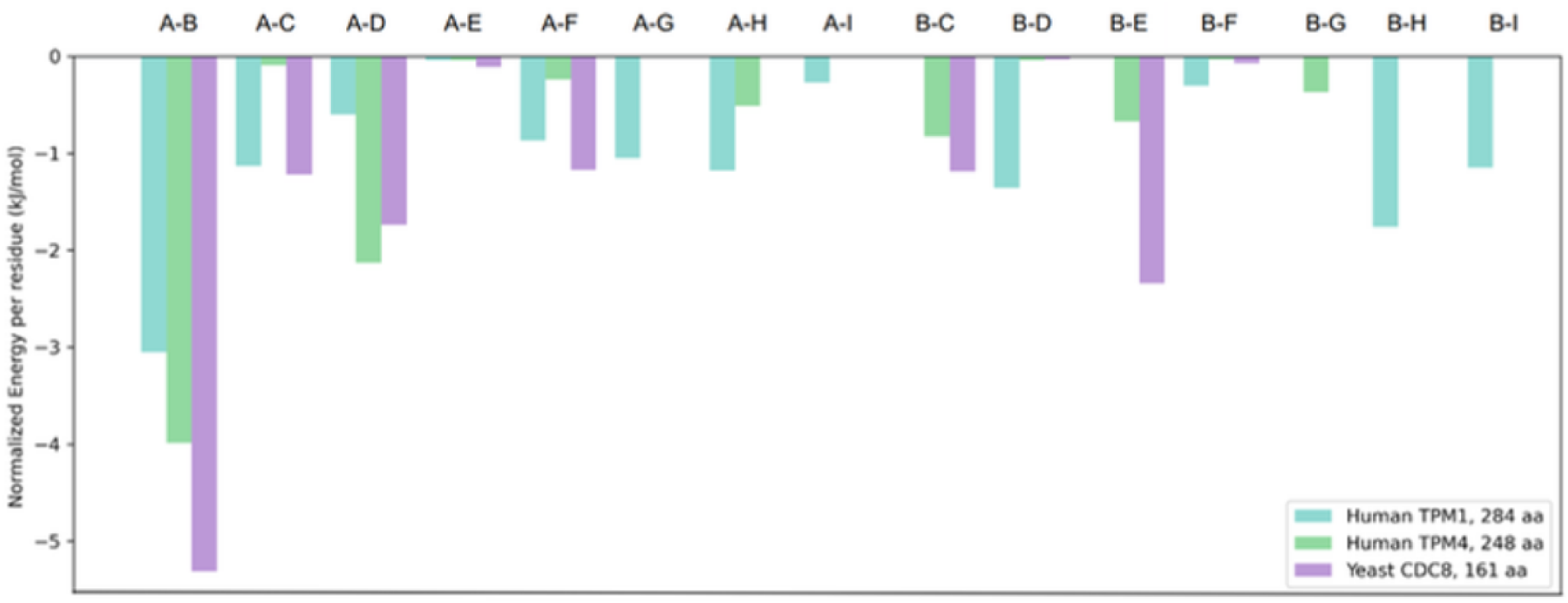
Normalized stabilizing energy per residue for each actin-tropomyosin interface in the docked complexes. Each group of bars corresponds to a tropomyosin-actin chain pair, indicated by the chain labels on the x-axis. In all complexes, chains A and B are the tropomyosin molecules, whereas chains C onwards are actin subunits; more negative values indicate more stabilizing interactions.

### Hotspot mapping

Hotspot residues were predicted using PPCheck. Despite sequence divergence, conserved hotspots were detected across equivalent actin-binding regions in all three tropomyosin molecules. As illustrated in Figure 4, an aspartate residue (D254 in hTPM1, D218 in hTPM4, and D131 in SpCdc8) occupies an equivalent position and is consistently classified as a hotspot. Similarly, E223 in hTPM1 (P6) and E187 in hTPM4 (P5) represent equivalent positions at which the same residue is conserved as a hotspot. Many of these hotspots are acidic and form salt-bridge interactions with the respective actin partners, highlighting the importance of conserved electrostatic contacts in stabilizing the interface. A complete list of hotspot residues is provided in Supplementary Table S3.

**Figure 4.**
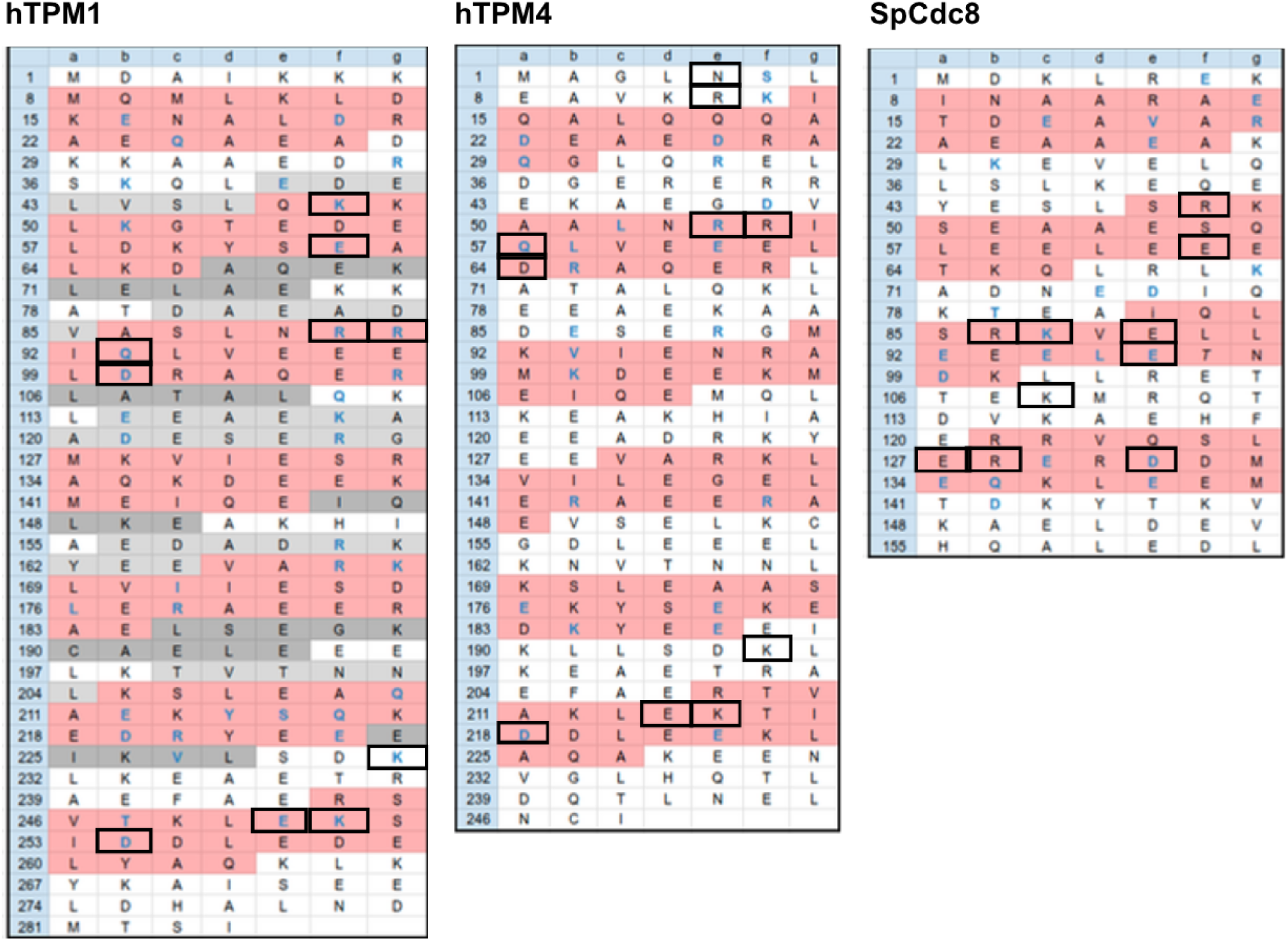
The sequences of hTPM1, hTPM4, and SpCdc8 split into heptads, with residues labelled a-g per heptad. Pink coloured cells represent actin binding regions adapted from <u>Mar</u>g<u>aret et al. 2012</u>. and cells with bolded blue text highlight hotspot residues identified using PPCheck. The residue number of each ‘a’ residue is shown on the left side. Cells with black borders represent hotspot residues observed in hTPM1 which are either fully or partly conserved in hTPM4 and SpCdc8 (example provided in text:: D254 in hTPM1, D218 in hTPM4, and D131 in SpCdcS).

### Temperature-sensitive *S. pombe* CDC8 mutants

Across mutants, the predicted stabilising energies were comparatively lower than that of the wild type SpCdc8 (Table 2), despite all coiled coils having approximately the same number of interface residues.

**Table 2.** Table displaying the total stabilising energies between the two tropomyosin chains in the coiled coil across SpCdc8 and known temperature-sensitive mutant variants. The data for the wild-type SpCdc8 is emphasised in bold font.

| Complex | Chain 1 | Chain 2 | Total Stabilizing Energy (kJ/mol) | Number of interface residues | Normalized Energy per residue (kJ/mol) |
| --- | --- | --- | --- | --- | --- |
| **SpCdc8** | **A** | **B** | **-1689.55** | **318** | **-5.31** |
| SpCdc8_A18T | A | B | -1077.89 | 315 | -3.42 |
| SpCdc8_E129K | A | B | -1155.1 | 315 | -3.67 |
| SpCdc8_E31K | A | B | -1154.24 | 314 | -3.68 |
| SpCdc8_R21H | A | B | -1055.79 | 315 | -3.35 |

Three of these mutations – A18T, R21H and E129K – occur in actin-binding regions (Figure 5A). However, there appeared to be no significant differences between the energies observed for actin-binding with wild type SpCdc8 and the four mutants (Table 3). A possible explanation is that the mutated residues and their neighbours do not interact with the actin subunits in the models (Figure 5B), on account of the distance between the residues being greater than 10 Å. These results are discussed further in the next section.

**Figure 5.**
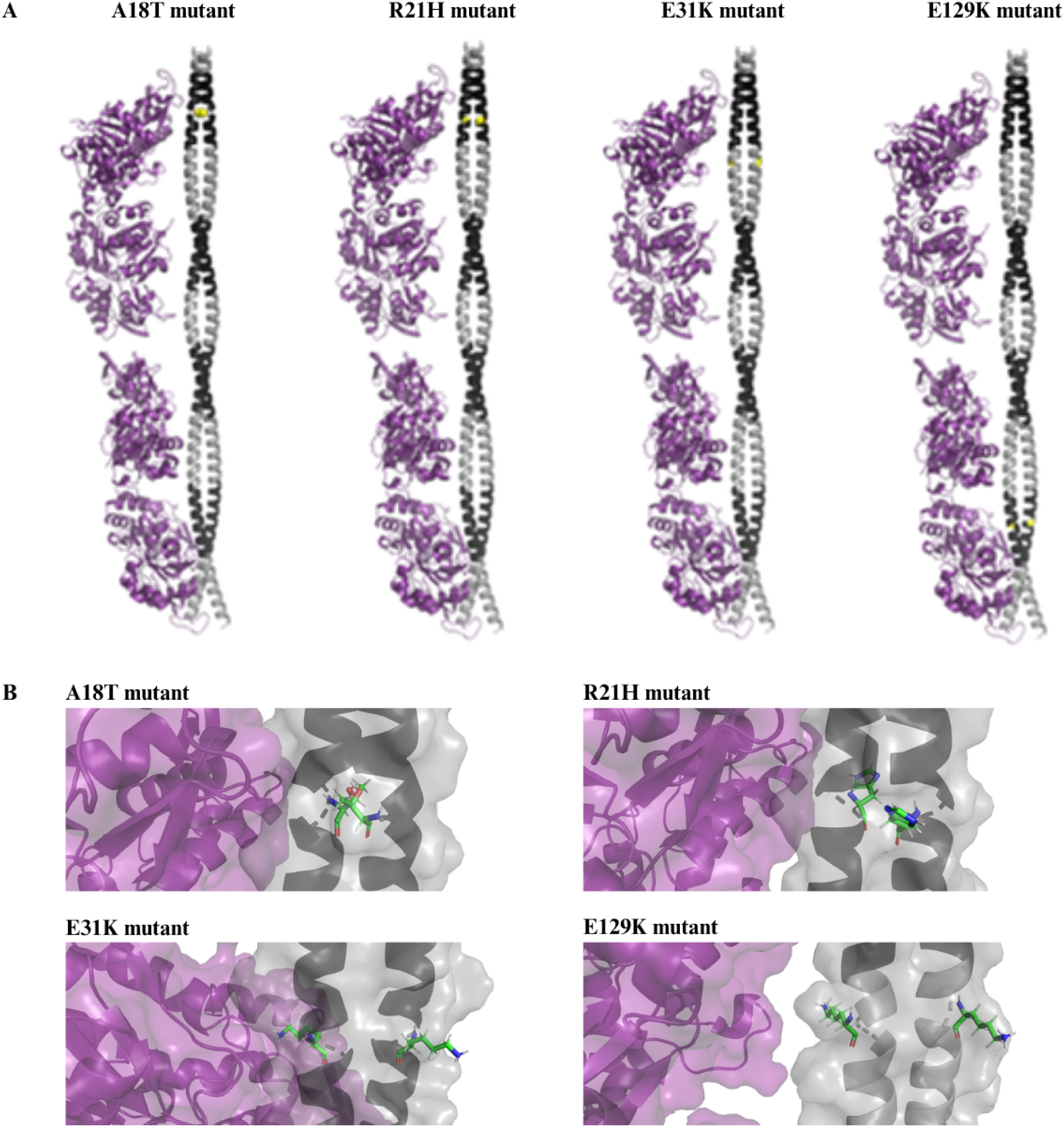
Modeled structures of SpCdc8-actin complexes with SpCdcS mutations (A18T, R21H, E31K, and E129K) that result in sensitivity to higher temperatures. The coiled-coiled is coloured grey, and actin-binding regions are indicated in black. Actin molecules are shown in purple. (A) Full length structures. The mutation sites are represented by yellow spheres. (B) Zoomed in view of the mutation sites, with the mutated residues shown as sticks. In all cases, the mutated residues are at distances of more than 10 angstrom from the corresponding actin molecule.

**Table 3.** Table displaying the total stabilising energies between the two tropomyosin chains (A and B) and the actin subunits (C-F) for the wild-type SpCdc8 and known temperature-sensitive mutant variants.

| Complex | Chain 1 | Chain 2 | Total Stabilizing Energy (kJ/mol) | Number of interface residues | Normalized Energy per residue (kJ/mol) |
| --- | --- | --- | --- | --- | --- |
| pCDC8-actin complex | A | C | -58.76 | 48 | -1.22 |
|  | A | D | -80.06 | 46 | -1.74 |
|  | A | E | -1.67 | 15 | -0.11 |
|  | A | F | -40.79 | 35 | -1.17 |
|  | B | C | -47.5 | 40 | -1.19 |
|  | B | D | -0.28 | 10 | -0.03 |
|  | B | E | -72.52 | 31 | -2.34 |
|  | B | F | -0.8 | 11 | -0.07 |
| pCDC8-A18T-actin complex | A | C | -51.84 | 41 | -1.26 |
|  | A | D | -40.78 | 37 | -1.1 |
|  | A | E | -1.07 | 11 | -0.1 |
|  | A | F | -30.86 | 29 | -1.06 |
|  | B | C | -19.54 | 30 | -0.65 |
|  | B | D | -0.06 | 2 | -0.03 |
|  | B | E | -9.83 | 25 | -0.39 |
|  | B | F | -0.7 | 10 | -0.07 |
| pCDC8-R21H-actin complex | A | C | -42.79 | 40 | -1.07 |
|  | A | D | -39.75 | 37 | -1.07 |
|  | A | E | -1.09 | 10 | -0.11 |
|  | A | F | -34.27 | 29 | -1.18 |
|  | B | C | -31.09 | 31 | -1 |
|  | B | D | -0.08 | 2 | -0.04 |
|  | B | E | -7.93 | 24 | -0.33 |
|  | B | F | -0.58 | 10 | -0.06 |
| pCDC8-E31K-actin complex | A | C | -49.24 | 45 | -1.09 |
|  | A | D | -37.38 | 37 | -1.01 |
|  | A | E | -0.96 | 11 | -0.09 |
|  | A | F | -33.15 | 31 | -1.07 |
|  | B | C | -38.93 | 30 | -1.3 |
|  | B | D | -0.1 | 2 | -0.05 |
|  | B | E | -7.93 | 23 | -0.34 |
|  | B | F | -0.49 | 10 | -0.05 |
| pCDC8-E129K-actin complex | A | C | -51.43 | 46 | -1.12 |
|  | A | D | -39.73 | 37 | -1.07 |
|  | A | E | -1.1 | 10 | -0.11 |
|  | A | F | -21.22 | 26 | -0.82 |
|  | B | C | -29.44 | 31 | -0.95 |
|  | B | D | -0.03 | 2 | -0.02 |
|  | B | E | -11.06 | 24 | -0.46 |
|  | B | F | -0.87 | 10 | -0.09 |

## Discussion

Our docking analysis suggests that filament length imposes a geometric constraint on actin affinity. Across the three isoforms examined, the shortest filament, *S. pombe* Cdc8, consistently formed the most stabilizing and residue-rich interface with actin, whereas the longer human isoforms showed weaker overall interface energies. This behavior is consistent with the “gestalt-binding” model (Holmes and Lehman, 2008), in which tropomyosin is viewed as a form-fitting, end-to-end polymer whose specificity arises from the concerted behavior of the entire acto–tropomyosin assembly rather than from a small set of high-affinity contacts. Over longer contour lengths, any mismatch between the intrinsic bending stiffness of tropomyosin (with a contour length of ∼40 nm and a persistence length of similar magnitude for isolated α-tropomyosin) and the geometry of the actin helix is expected to accumulate as angular deviations, which can locally weaken binding (Loong et al., 2012). In our models, the shorter SpCdc8 filament minimizes these cumulative deviations along the actin groove and thereby supports a tighter, more side-on continuous interface, whereas extended human filaments incur a greater steric and elastic penalty for maintaining registry over multiple actin subunits. Repeated LightDock runs for hTPM1 consistently converged on a superhelical trajectory compatible with the native actin helix, indicating that the observed superhelical arrangement arises naturally from the docking procedure rather than from imposed restraints.

Within the human isoforms, our computational results are consistent with the functional classification emerging from isoform-resolved experimental studies. The muscle-associated hTPM1 exhibits more stabilizing actin contacts and a higher density of interface residues in our docking models, mirroring a well-established distinction between high-molecular-weight (HMW) and low-molecular-weight (LMW) tropomyosins. HMW isoforms, including sarcomeric TPM1 products, are enriched in contractile structures that demand robust, cooperative activation, whereas LMW isoforms, including many TPM4 products, are preferentially localized to dynamic cytoskeletal populations such as stress fibers and cell edges (Rao et al., 2012; Schevzov et al., 2011; Vindin and Gunning, 2013). In motility and reconstitution assays, HMW tropomyosins generally support stronger myosin-based motility and more stable actin filaments than LMW isoforms, which are instead associated with rapid turnover and remodeling (Barua et al., 2014; Creed et al., 2011). This difference has been directly captured by FRAP experiments examining tropomyosin exchange kinetics on actin filaments in living cells. Tojkander and colleagues compared GFP/YFP-tagged isoforms on ventral stress fibers of U2OS osteosarcoma cells using double-exponential fits of 10–20 recovery curves per isoform, resolving fast and slow exchange components. Both Tpm1.6 (Tm2) and Tpm1.7 (Tm3), which are the principal TPM1 gene products in non-muscle cells, exhibited substantially slower exchange kinetics than Tpm4.2 (Tm4): the slow-component halftimes for Tpm1.6 and Tpm1.7 were 161.2 ± 37.7 s and 113.6 ± 10.5 s, respectively, compared to 56.8 ± 13.2 s for Tpm4.2, indicating an approximately two-fold difference in the residence time of the stable bound population. The fast-component halftime of Tpm4.2 (4.8 ± 2.0 s) was similarly shorter than that of either Tpm1.6 (13 ± 2.5 s) or Tpm1.7 (9.4 ± 1.8 s) (Tojkander et al., 2011). Gateva and coworkers extended the comparison to purified proteins on reconstituted actin filaments by *in vitro* TIRF-FRAP, demonstrating that sfGFP-Tpm1.6 and sfGFP-Tpm1.7 displayed only very slow fluorescence recovery over a 10-minute post-bleach window, while sfGFP-Tpm4.2 showed rapid recovery on actin filament bundles, directly attributing this difference to the higher cooperativity of actin binding by HMW isoforms (Gateva et al., 2017). Together, these kinetic data establish that the slower exchange of TPM1 isoforms from actin filaments reflects a fundamentally higher actin-binding affinity relative to Tpm4.2, consistent with its deployment in cytoskeletal contexts requiring continuous filament remodeling. The stable actin interface predicted for hTPM1 in our docking models is therefore in agreement with its experimentally established role in cardiac and skeletal muscle thin filaments, while the comparatively weaker hTPM4 interface aligns with the rapid exchange kinetics observed for Tpm4.2 in cytoskeletal contexts requiring continuous filament re-modeling.

At the residue level, the three isoforms achieve these distinct geometric and functional outcomes while preserving a conserved set of energetic hotspots anchored by acidic residues. Our models identify conserved aspartate and glutamate residues (e.g. D254 in hTPM1, D218 in hTPM4 and the equivalent D131 in SpCdc8) that repeatedly form salt bridges to actin, together with additional conserved glutamates in equivalent positions of the quasi-repeats. These features are consistent with sequence and mutational analyses showing that evolutionarily conserved acidic residues in surface-exposed positions, especially within periods 2 and 5, make disproportionate contributions to actin affinity and cooperative activation (Barua et al., 2013). Studies have highlighted period 2 as a hotspot for electrostatically favorable actin–tropomyosin contacts in integrative models of the cardiac thin filament (Sunitha et al., 2012), while independent mutagenesis studies found that substitutions in the first half of periods 2, 4 and 5 strongly reduce actin binding (Barua et al., 2011). Our observation regarding the conservation of acidic hotspots across species and isoforms suggests that the actin–tropomyosin interface is built around a small number of important, highly conserved, electrostatic anchor points, onto which isoform-specific sequence changes then modulate local packing, as well as long-range filament mechanics.

A number of germline TPM4 variants with clear clinical relevance have been reported to date, most notably the non-sense mutations R69 (period 2) and Q108 (period 3), as well as the missense substitution R146C (period 4), all associated with platelet-type bleeding disorders (ClinVar). By contrast, TPM1 harbours a large and well-characterized spectrum of germline missense and truncating variants associated with inherited cardiomyopathies, including hypertrophic, dilated, and left ventricular non-compaction phenotypes (Gunning et al., 2008; Rao et al., 2012). These pathogenic variants are distributed along the entire length of the TPM1 coiled coil, with notable enrichment in the N-terminal overlap region (period 1; e.g. M8R, D14Y, K15N, E16Q), the central actin-interacting region (period 2; e.g. E40K, E54K, D55N, E62Q, A63V), and the C-terminal regulatory region (e.g. D175N and E180G in period 5, E192K between periods 5 and 6, and K248E in period 7) (Li et al., 2011; Sunitha et al., 2012). Several of these substitutions have been experimentally shown to reduce actin affinity or alter calcium-dependent regulation of myosin binding (Barua et al., 2013, 2011), underscoring the stringent structural and energetic constraints imposed on TPM1 in muscle thin filaments. The breadth and positional diversity of TPM1 disease-linked variants therefore stand in contrast to the comparatively sparse genetic landscape of TPM4, reinforcing the inference that muscle tropomyosins operate under tighter geometric and regulatory constraints than their cytoskeletal counterparts.

The behavior of *S. pombe* Cdc8 temperature-sensitive mutants further underscores the importance of tropomyosin filament mechanics rather than local actin contacts *per se*. Of these, the E31K mutation occurs in between periods 1 and 2; E129K is within period 4; A18T and R21H are present within period 2. In our models corresponding to A18T, R21H, E31K and E129K, the tropomyosin filaments maintain near-wild-type actin interface energies, but show reduced stability within the coiled-coil dimer, particularly at sites that influence N-terminal overlap and longitudinal continuity of the tropomyosin cable. Experimentally, N-terminal acetylation of Cdc8 has been shown to enhance actin affinity and proper localization, while temperature-sensitive cdc8 alleles (including cdc8-27 and cdc8-110) primarily disrupt actin cable integrity and contractile-ring assembly rather than abolishing actin binding outright (Christensen et al., 2019; Cranz-Mileva et al., 2013; Palani et al., 2020; Skoumpla et al., 2007). These *in vivo* data support a model in which such mutations reduce the thermal stability and end-to-end polymerization competence of the Cdc8 cable, thereby compromising the cooperative behavior of the thin filament at restrictive temperatures. In this view, regulatory failure arises not from a simple loss of local affinity but from a breakdown in the ability of the tropomyosin polymer to transmit conformational information and maintain continuous coverage of the actin filament.

Taken together, our results support a hierarchical view of tropomyosin–actin recognition. At the global scale, filament length and bending stiffness constrain how well a given tropomyosin can track the actin helix, with shorter filaments like Cdc8 incurring smaller cumulative geometric penalties and therefore achieving higher overall interface stability. At the intermediate, isoform level, tissue-specific demands on cooperativity versus plasticity are met by tuning the balance between stabilizing contacts and geometric frustration, exemplified by the stronger interfaces of muscle hTPM1 relative to cytoskeletal hTPM4. Finally, at the local scale, conserved acidic hotspots provide electrostatic anchor points that are preserved across species and regulatory states, while more peripheral residues and coiled-coil stability determinants accommodate evolutionary experiments such as temperature-sensitive mutations in Cdc8. These multi-scale design principles offer a structural rationale for how a single coiled-coil scaffold can be re-purposed across tissues and organisms to generate robustly cooperative or dynamic actin filaments.

## Conclusions

We conclude that tropomyosin regulation is governed by a balance of filament geometry and sequence-specific electrostatics. While shorter filaments like CDC8 inherently favour tighter packing against the actin helix, human isoforms have evolved specific interfacial chemistries to tune this affinity, resulting in strong binding for muscle isoforms (TPM1) and dynamic binding for cytoskeletal isoforms (TPM4). These structural insights, supported by the energetic hierarchies established in previous literature, provide a unified model for how tropomyosin length and sequence determine its stability.

