## SupplementalFigures for "Comparative Modelling of Actin–Tropomyosin Interfaces"

### Figure S1

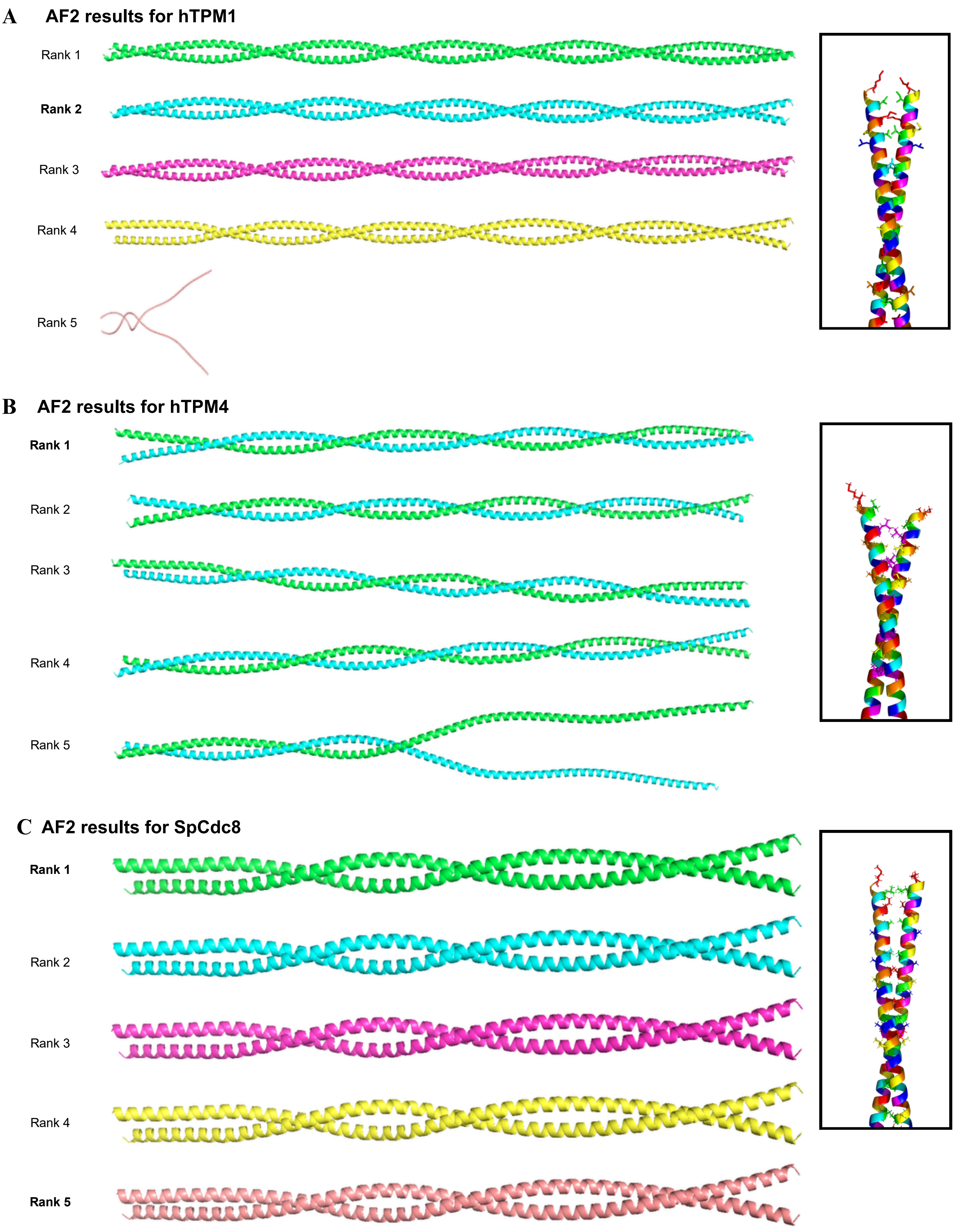


**Figure S1.** Alphafold2-generated structures for (A) hTPM1, (B) hTPM4, and (C) SpCdc8. Insets show a zoomed-in view of part of the dimers selected for iterative docking. Dimers are coloured according to the *abcdefg* heptad repeat to illustrate the dimerization interface. The hydrophobic core positions, **a** (red) and **d** (green), interlock at the dimer interface. This core is flanked by positions **e** (cyan) and **g** (magenta), which often participate in inter-helical electrostatic interactions and salt bridges. Solvent-exposed positions **b**, **c**, and **f** are colored orange, yellow, and blue, respectively. Sidechains are shown as sticks; all structures visualized using PyMOL.

### Figure S2

#### A CoCoNat results

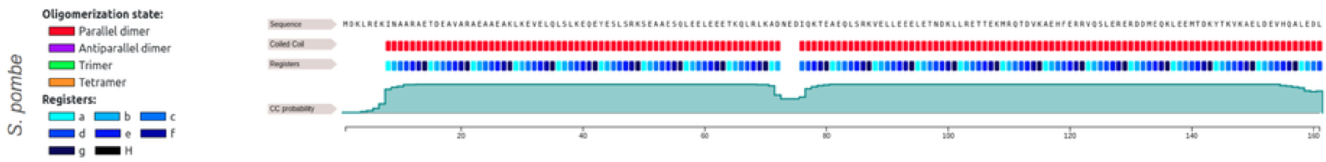


#### DeepCoil2 results

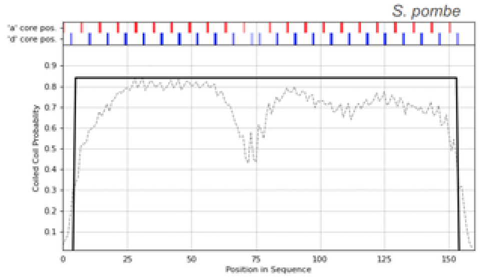


### B

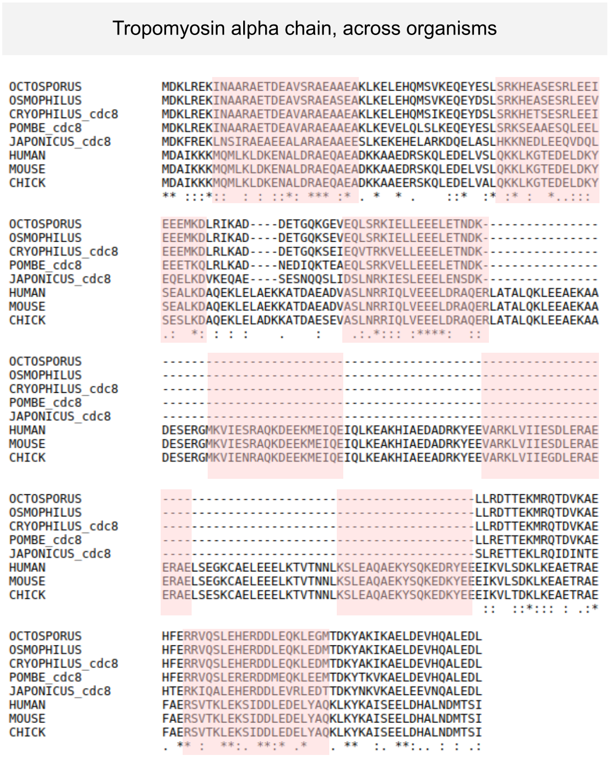


### C

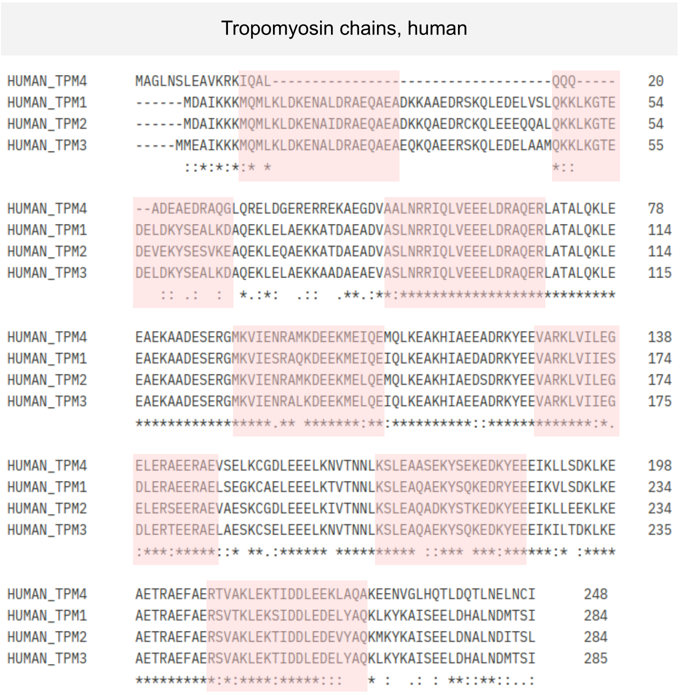


**Figure S2.** (A) DeepCoil2 and CoCoNat results for SpCdc8 validating its coiled-coiled structure with heptad registry. (B) Sequence alignments of tropomyosin alpha chain with other vertebrate and yeast homologues. (C) Sequence alignment of human tropomyosin isoforms. Both sets of sequence alignments were performed using ClustalW, and the pink blocks represent actin binding regions adapted from Margaret et al. 2012,
